# Ecological specificity of the metagenome in a set of lower termite species supports contribution of the microbiome to adaptation of the host

**DOI:** 10.1101/526038

**Authors:** Lena Waidele, Judith Korb, Christian R Voolstra, Franck Dedeine, Fabian Staubach

## Abstract

**Background:** Elucidating the interplay between hosts and their microbiomes in ecological adaptation has become a central theme in evolutionary biology. A textbook example of microbiome-mediated adaptation is the adaptation of lower termites to a wood-based diet, as they depend on their gut microbiome to digest wood. Lower termites have further adapted to different life types. Termites of the wood-dwelling life type never leave their nests and feed on a uniform diet. Termites of the foraging life type forage for food outside the nest and have access to other nutrients. Here we sought to investigate whether the microbiome that is involved in food substrate breakdown and nutrient acquisition might contribute to adaptation to these ecological differences. We reasoned that this should leave ecological imprints on the microbiome.

**Results:** We investigated the protist and bacterial microbiomes of a total of 29 replicate colonies from five termite species, covering both life types, using metagenomic shotgun sequencing. The microbiome of wood-dwelling species with a uniform wood diet was enriched for genes involved in lignocellulose degradation. Furthermore, metagenomic patterns suggest that the microbiome of wood-dwelling species relied primarily on direct fixation of atmospheric nitrogen, while the microbiome of foraging species entailed the necessary pathways to utilize nitrogen in the form of nitrate for example from soil.

**Conclusion:** Our findings are consistent with the notion that the microbiome of wood-dwelling species bears an imprint of its specialization on degrading a uniform wood diet, while the microbiome of the foraging species might reflect its adaption to access growth limiting nutrients from more diverse sources. This supports the idea that specific subsets of functions encoded by the microbiome can contribute to host adaptation.

## Background

The importance of microbes for the evolution of higher organisms is starting to be realized [1, 2]. Metazoan evolution is not only driven by pathogenic microbes, as reflected by fast evolution of immune genes [3]. Rather, microbes often are facilitators of metabolic and environmental adaptations [2, 4, 5]. For instance, the gut microbial communities of wood-feeding roaches and termites facilitate thriving on a wood diet that is difficult to digest and poor in nitrogen. Nitrogen fixation and the digestion of wood depend on the termite gut microbiome [2, 6, 7]. In lower termites, lignocellulose degradation was initially mainly attributed to unicellular eukaryotes (protists) in the gut [8]. Recently, it has become evident that lignocellulose degradation is a synergistic effort of the termite, its associated protists, and bacteria [9–11]. In addition to their role in lignocellulose degradation, bacteria are also essential for the assimilation of nitrogen taken up from the environment. Nitrogen can be acquired from the environment either via fixation from the atmosphere [12, 13], or via nitrate reduction [14]. Also, nitrogen can be recycled from the metabolic waste product uric acid [15, 16]. By using genome sequencing and pathway reconstruction, these processes have been assigned to four major bacterial phyla in the termite gut: Proteobacteria (*Desulfovibrio* [17]), Spirochetes (*Treponema* [18, 19]), Bacteroidetes (*Azobacteroides*) [16], and Elusimicrobia (*Endomicrobium* [20, 21]).

Many bacteria in the termite gut live in tight association with protists, where they sit on the surface [22, 23], in invaginations of the cell membrane [17], or even inside the protist cells [24]. Such tight associations lead to frequent vertical transmission of bacteria between protist generations. In return, protists and bacteria are vertically transmitted between termite generations via proctodeal trophallaxis during colony foundation [25]. Vertical transmission has led to co-speciation between bacteria and their protist hosts, and sometimes even the termite hosts [26–29]. Evidence for horizontal transfer of protists between termite species, so called transfaunations, is limited to a few exceptions [30]. Hence, the termite host species association is rather strict, leading to strong phylogenetic imprints on protist community structure [31–33]. In comparison, the bacterial microbiome is more flexible, frequently transferred between termite host species [34], and affected by diet [33, 35–41].

There is evidence that the gut microbiome of termites has contributed to the adaptation of different termite species to their specific ecologies [33, 36, 42–44]. There are pronounced ecological differences between the so-called termite life types [45, 46]. Termite species of the wood-dwelling life type never leave their nest except for the mating flight. They feed on a relatively uniform bonanza resource, that is the piece of wood they built their nest in [47, 48]. On the other hand, foraging species leave their nest to forage for food and have access to additional nutrients [47, 49]. This likely imposes different selection pressures on the termite holobiont, in particular with regard to nutrient uptake. Because the microbiome is directly involved in nutrient uptake, it seems reasonable to hypothesize that it may also play a role in adaptation to life type related ecological differences. In this scenario, one would expect the life types to leave an imprint on microbiome structure and function. As such, searching for microbial imprints of a given life type can possibly provide us with a lead for microbiome-mediated adaptation.

One potential pitfall of such an endeavor is that microbiomes may bear imprints from transient microbes that were ingested from the environment. Transient microbes rarely form evolutionary relevant relationships with the host [50, 51]. Instead, they reflect short-term associations with microbes from the local environment the termites were collected from. When the local environment correlates with other ecological differences, these imprints could be falsely interpreted as potentially adaptive changes of the microbiome. Therefore, it is essential to reduce the impact of these microbes on the analysis. In order to explore potential ecological imprints on the microbiome, we focused on an evolutionary switch between wood-dwelling and foraging life types in the Rhinotermitidae (Figure 1). *Reticulitermes* species are of the foraging life type, while *Prorhinotermes simplex* is wood-dwelling. If the microbiome was affected by life type specific ecology, we would expect that the microbiome of *Prorhinotermes simplex* was similar to that of the other wood-dwelling species (*Cryptotermes*) although these are from a different family (Kalotermitidae). At the same time, the microbiome of the foraging *Reticulitermes* species should bear distinct features. Alternatively, if there was no ecological imprint, we would expect the microbiome to follow a phylogenetic pattern, with the Rhinotermitidae *Prorhinotermes* and *Reticulitermes* forming a cluster and the *Cryptotermes* species (Kalotermitidae) forming a second cluster. Using this experimental setup, we recently showed that protist community composition aligned with phylogeny, but bacterial communities aligned more strongly with wood-dwelling and foraging life types [33].

**Figure 1:**
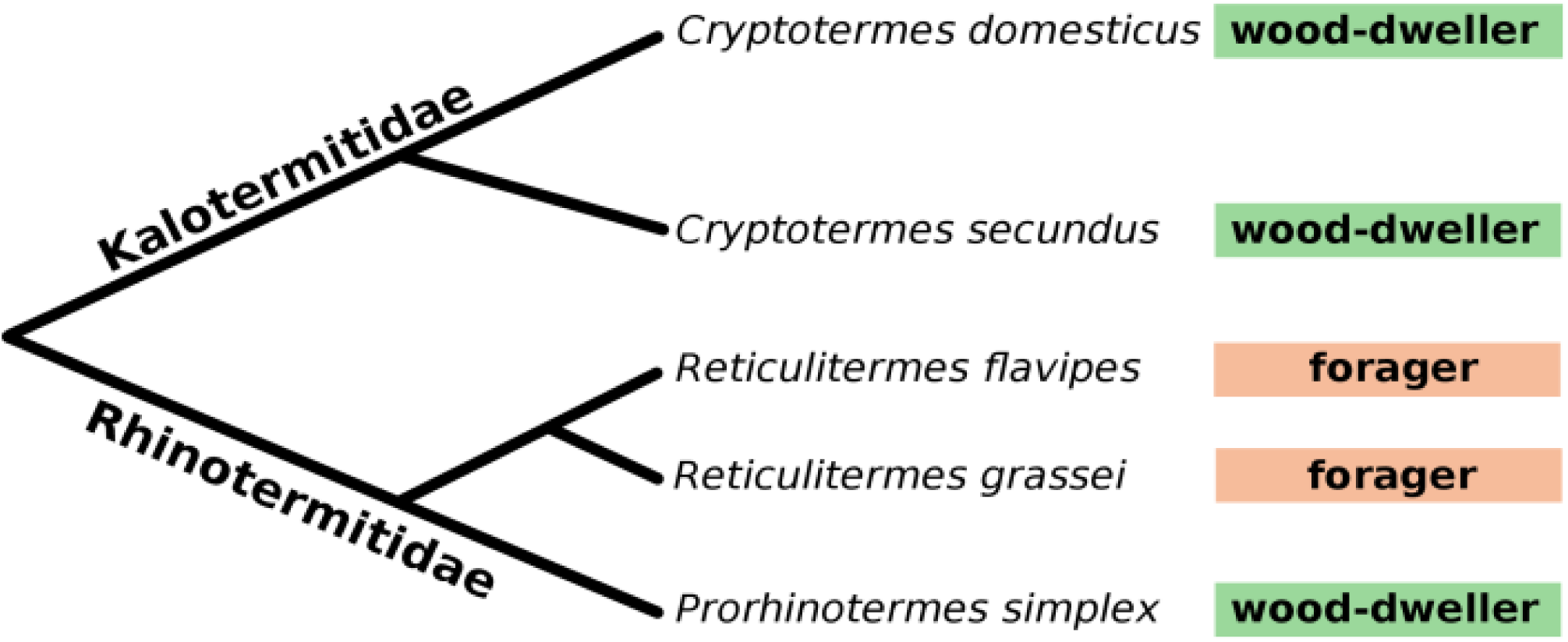
Schematic phylogeny of the five lower termite species used in this study from [33]. Branch length not drawn to scale. Colored boxes indicate the life type.

To further explore this, we investigated whether changes in microbiome composition are also reflected by changes in microbiome function, as would be expected if the microbiome played a role in adaptation. For instance, we would expect dietary adaptations to be reflected by changes to pathways involved in substrate breakdown and effective provisioning of limiting nutrients such as nitrogen. In order to test whether and which changes in the functional repertoire align with life type and could be involved in potential adaptation to different ecologies, we characterized the metagenome of two foraging species; *Reticulitermes flavipes* and *Reticulitermes grassei.* We compared their functional repertoire to that of three wood-dwelling species *Prorhinotermes simplex*, *Cryptotermes secundus*, and *Cryptotermes domesticus.* Because there can be substantial variation in microbial communities between colonies [52–55], we analyzed five *C. domesticus*, eight *C. secundus*, seven *P. simplex*, five *R. flavipes*, and four *R. grassei* replicate colonies. We focused on the persistent, long-term differences between microbiomes by controlling short-term effects caused by the influx of transient microbes. This was achieved by feeding a common diet of sterile *Pinus* wood for several weeks prior to sample collection.

## Results

We analyzed a total of ∼440 million metagenomic shotgun sequences. Between 974,176 and 8,949,734 sequences per sample were of microbial origin (Table S1). Sequences were subsampled (rarefied) to 1,386,882 bacterial and 2,781 protist annotated sequences per sample. For annotation, sequences were aligned to a reference database of clusters of orthologous groups of genes (COGs) with known function. These COGs represent the lowest level of the eggNOG hierarchical annotation. At the next higher level, the COGs are grouped into pathways (Figure S1, Figure S9), and at the third and highest level, the pathways are grouped into three categories; “information storage and processing”, “cellular process and signaling”, and “metabolism”. We adhere to this definition of the eggNOG hierarchical terms throughout the study.

“information storage and processing” differentiates the protist metagenomes of wood-dwelling and foraging lower termite species

In our previous study [33] on the identical samples, the protist communities of the Rhinotermitidae *Prorhinotermes* and *Reticulitermes* clustered together, supporting a phylogenetic imprint on community composition. Here, we tested whether this pattern was also reflected by the functions encoded by the protist metagenome. Therefore, we annotated metagenome encoded functions in the shot gun sequences and compared the functional metagenome profiles across host species, using Bray-Curtis-Dissimilarity [56]. This index considers abundance of functional categories, thus avoiding arbitrary coverage cutoffs.

The protist functional repertoire clustered according to host family and genus (Figure 2A), thus showing a dominant phylogenetic imprint. Family wise clustering was supported by Redundancy Analysis (RDA): the model including host family explained more variance in the functional repertoire and produced lower AICs than the model based on life type (Table 1). For a more detailed view, we analyzed the three categories at the highest level in the eggNOG hierarchical annotation (Figure S1) separately. Cluster analysis of the categories “cellular process and signaling” and “metabolism” supported the notion that phylogenetic relatedness is an important factor for functional similarity (Supplement Figures S4B and D). In contrast, the portion of the metagenome assigned to “information storage and processing” (Figure 2B) clustered primarily by life type. The stronger effect of life type than phylogeny on this functional category was also supported by higher explanatory power and lower AICs in RDA (Table 1).

**Figure 2:**
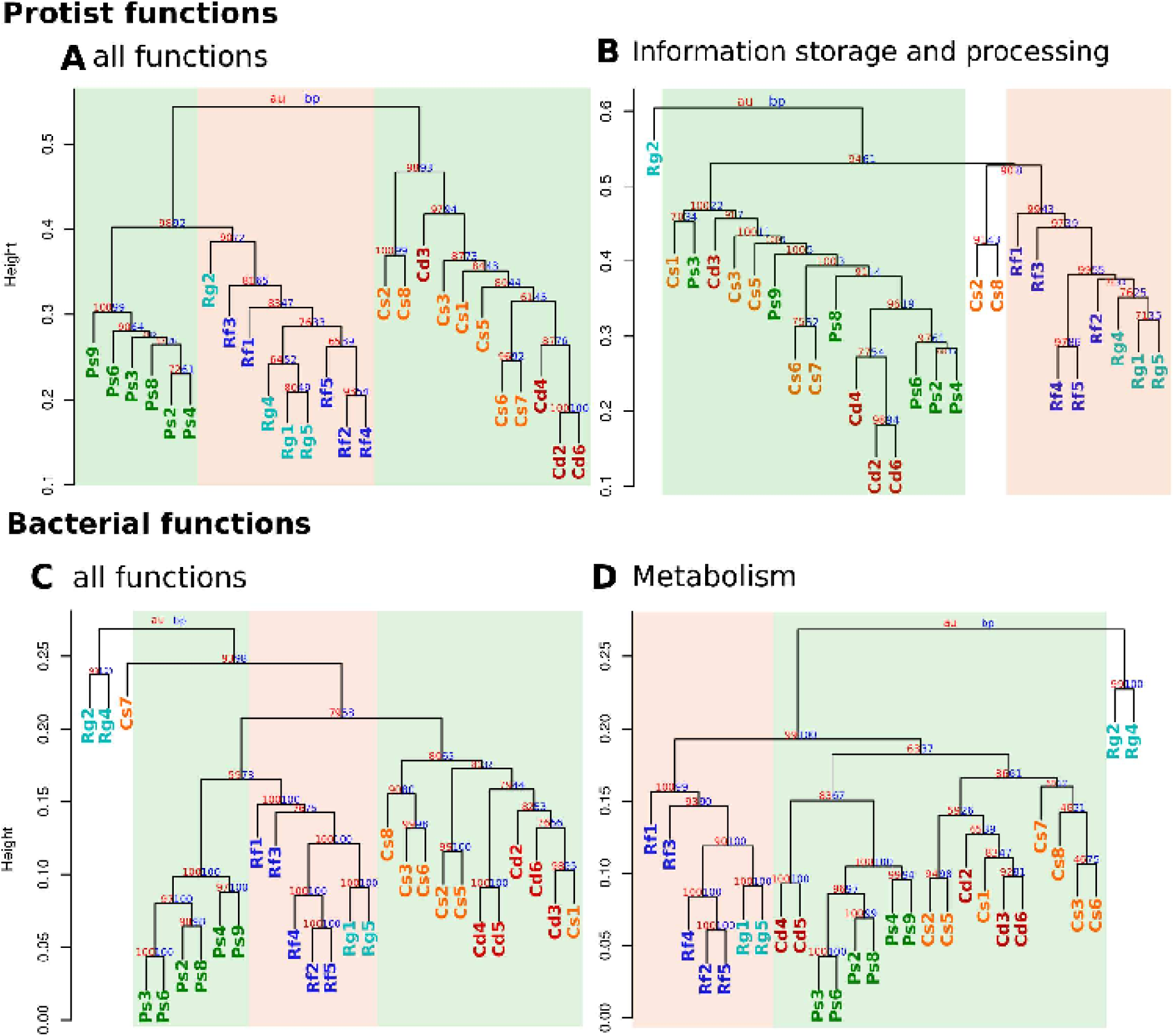
Cluster dendrograms of the functional profiles of the protist and bacterial community. Community distances are based on Bray-Curtis Dissimilarities of A) all functions of the protist community (25,795 sequences), B) The category “information storage and processing” of the protist community (4,527 sequences), C) all functions of the bacterial community (21,215,480 sequences), and D) the category “metabolism” of the bacterial community (10,586,058 sequences). Cd (red) = *C. domesticus* colonies; Cs (orange) = *C. secundus* colonies; Ps (green) = *P. simplex* colonies; Rf (blue) = *R. flavipes* colonies; Rg (lightblue) = *R. grassei* colonies. Green background = wood-dwelling life types; orange background = foraging life type. For protist functions involved in “information storage and processing” in the protist community, samples clustered according to life type. Similarly, the bacterial metabolic metagenomes clustered according to life type. Cluster dendrograms of all functional categories of protist and bacterial communities can be found in Supplementary Figure S4 and S5.

**Table 1:** Models of the effects of life type and host family (phylogeny) on functional community profiles. Effects were analyzed with Redundancy analysis using the functional abundance table as response variable. Host family was the best explanatory variable for the combination of all protist functions, however life type explained more variance (R²) and produced a lower AIC for the category “information storage and processing” than host family. For all bacterial functions taken together, host family again explained a larger proportion of the variance, while life type was the best explanatory variable in the category “metabolism”.

| | | model | AIC | $R^2$ | $p$ |
| --- | --- | --- | --- | --- | --- |
| **protists** | all functions | null | -41.8 |  |  |
|  |  | host family | -49.0 | 0.27 | 0.001 |
|  |  | host life type | -45.4 | 0.16 | 0.001 |
|  | information storage and processing | null | -36.8 |  |  |
|  |  | host life type | -39.0 | 0.11 | 0.001 |
|  |  | host family | -38.4 | 0.09 | 0.001 |
| **bacteria** | all functions | null | -85.0 |  |  |
|  |  | host family | -92.7 | 0.27 | 0.001 |
|  |  | host life type | -90.1 | 0.20 | 0.001 |
|  | metabolism | null | -92.4 |  |  |
|  |  | host life type | -98.1 | 0.22 | 0.001 |
|  |  | host family | -96.8 | 0.18 | 0.001 |

Identification of the functions that differentiate the protist metagenomes of the wood-dwelling and foraging species can hold clues with regard to the nature of potentially adaptive phenotypes in the protist metagenome. In order to do so, we performed a linear discriminant analysis (LEfSe: [57]). This analysis identified 22 over-represented COGs in foraging and 14 in wood-dwelling species (Figure 3A, Table S3, p < 0.05, q < 0.05, LDA score > 2, Figure S6).

**Figure 3:**
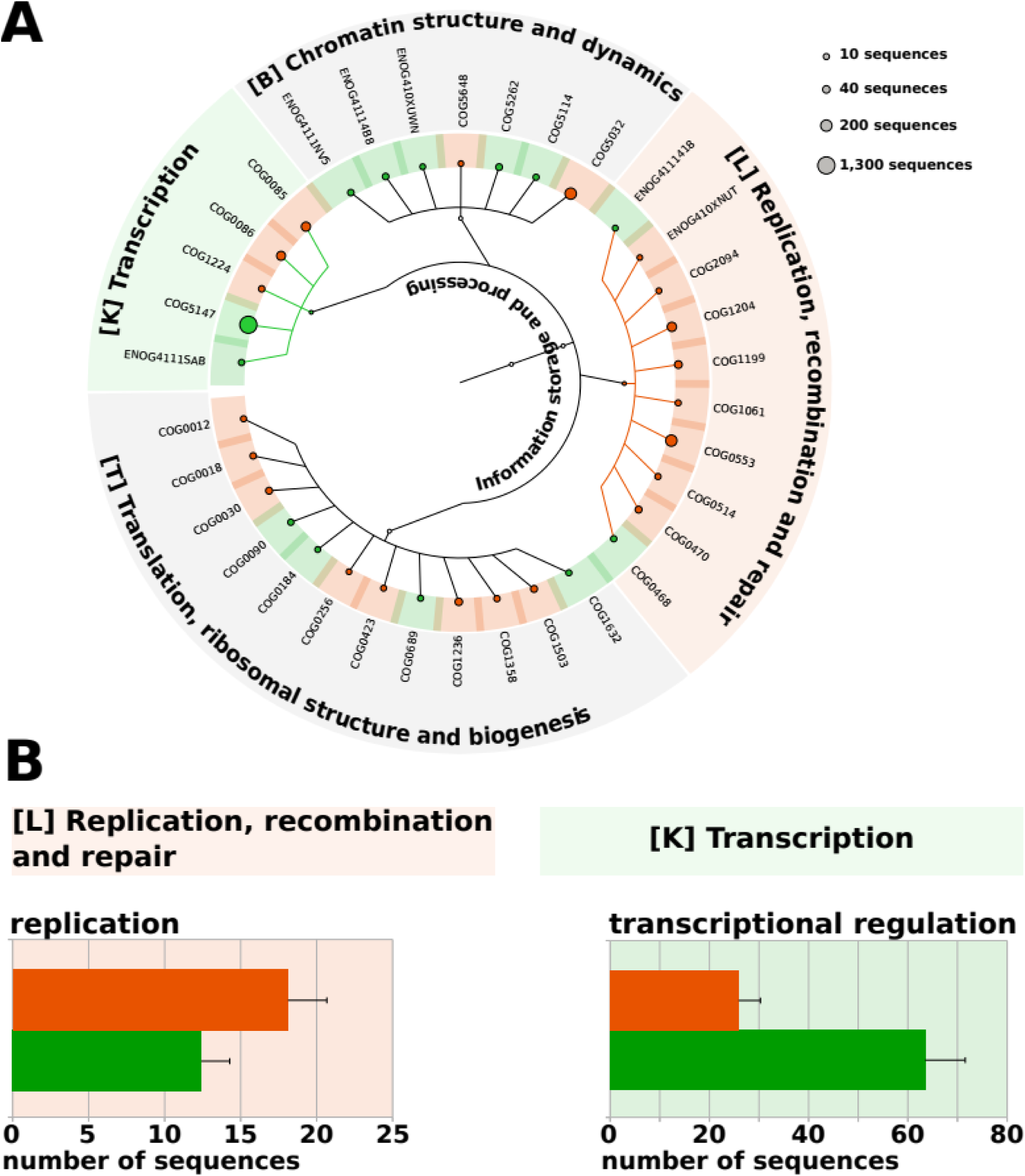
Differences in the functional content of the protist metagenomes of wood-dwelling and foraging species. A) Circular dendrogram/hierarchy of all over-represented COGs in the category “information storage and processing” in wood-dwelling species (green) or foraging species (orange). Circle size at the edges scales with abundance of the COG. Colored branches indicate over-represented pathways. Over-representation was detected with LEfSe [57] (p < 0.05, q < 0.05, LDA > 2). A Venn diagram visualizing the total number, and the differentially abundant number of functions in each of the five pathways that constitute the category “information storage and processing” can be found in Supplementary Figure S6. B) Sequence coverage of wood-dwelling (green) and foraging (orange) species of examples of over-represented COGs mentioned in the text. Error bars represent 95% confidence intervals across replicate colonies.

The pathway “replication, recombination, and repair” was over-represented in the foraging species (Figure 3A, Table S3, p = 0.0001, q = 0.002). The over-represented COGs in this pathway included a DNA dependent DNA-polymerase (COG0470) and five helicases (COG0514, COG0553, COG1199, COG1204, ENOG410XNUT, see Figure 3B for grouped analysis and Table S3 for individual COG p- and q-values). In wood-dwelling species, the pathway “transcription” was over-represented (p = 0.0004, q = 0.003). The over-represented COGs in this pathway contained DNA binding domains and were supposedly involved in transcriptional regulation (COG5147, ENOG4111SAB).

### The bacterial metabolic metagenome aligns with host ecology

In our previous study [33], the bacterial community composition of termite hosts clustered primarily by life type, which is consistent with ecology related differences between microbiomes. Following the rationale above, we tested whether this pattern was also reflected by the functions encoded by the metagenome.

Against the expectation from our previous study, the functional bacterial profiles showed no life type, but a phylogenetic imprint, which is in line with the protist functional profiles. Most samples clustered according to host family (Figure 2C). Analyzing the three high-level eggNOG functional categories separately provided more detailed insight. The categories “cellular process and signaling” and “information storage and processing” supported the notion of strong phylogenetic effects on metagenome function (Supplement Figures S5B and C). In contrast, the metabolic metagenomes (Figure 2D) clustered primarily according to host life type. Host life type was also a better predictor for metabolic functions than host family in RDA (Table 1).

Aside from these general patterns, several samples stood out. Samples Rg2 and Rg4 of *R. grassei* were on long branches in the dendrograms (Figure 2 and Supplement Figure S5), suggesting unusual functional profiles. Notably, these samples already stood out in our previous study [33] because of their unusual abundance of microbial taxa potentially due to infection with pathogens. This unusual composition was confirmed by taxonomic annotation in this study (see Supplement Figure S3). Sample Cs7 (*C. secundus*) also clustered separately from the other samples. This was mainly driven by abundant transposases in this sample (53.1% of sequences) (for example COG1662, COG3385, or ENOG410XT1T, see Table S2), accompanied by an increase in the frequency of *Bacteroides* (Figure S3) that are rich in conjugative transposons [58, 59]. We performed all analyses with and without these samples and found no qualitative differences (data not shown).

Bacterial metabolic functions that differentiated wood-dwelling from foraging species were identified using linear discriminant analysis (LEfSe). 105 metabolic COGs were over-represented in the wood-dwelling species, while 151 were over-represented in the foraging species (Table S4, p < 0.05, q < 0.05, LDA score > 2, Figure S7). All COGs described as over-represented or enriched in the following were subject to these p-value, q-value and LDA cutoffs. Because of their specialized diet, genes involved in nitrogen metabolism and lignocellulose break down like gycoside hydrolases (GH) are of particular interest, when focusing on metabolic differences among gut microbiomes in wood-feeding termites with different ecologies. In fact, among the genes involved in ‘carbohydrate transport and metabolism’ that were enriched in the microbiome of wood-dwelling termites, GHs were over-represented (43.3% of enriched genes versus 12% expected, exact binomial test: p = 2.124e-05, Table S4, S5). In the foraging termite species, only one gene with putative lignocellulolytic activity was over-represented (COG3858), suggesting that the wood-dwelling species have a higher potential for complex carbohydrate degradation. To further investigate differences in GH abundance between the microbiomes of wood-dwelling and foraging species, we performed a detailed pathway analysis using the CAZy database ([60], Figure 4). All GHs acting in hemicellulose break-down were more abundant in the wood-dwelling species (Figure 4B). Among the cellulolytic enzymes, ß-glucosidases were significantly more abundant in the wood-dwelling species. The other two enzymes involved (cellulase (endo-ß-1.4-glucanase), cellobiohydrolase) showed a trend into the same direction. All of the genes with cellulolytic or hemicelllulolytic activity were affiliated with Bacteroidetes (mostly members of the genus *Bacteroides*) or the genus *Treponema*. Additional support for the increased importance of hemicellulose utilization in the wood-dwelling species, was provided by the over-representation of twelve COGs annotated as TonB-dependent receptors (ENOG410XNNV, ENOG410XNPQ or COG4206, see Table S4). Apart from other substrates, these receptors are important for the uptake of plant-derived hemicellulose [61, 62]. All functions annotated as TonB-dependent receptors (or TonB-dependent associated receptor plugs) were affiliated with the genus *Bacteroides* (see Table S4).

**Figure 4:**
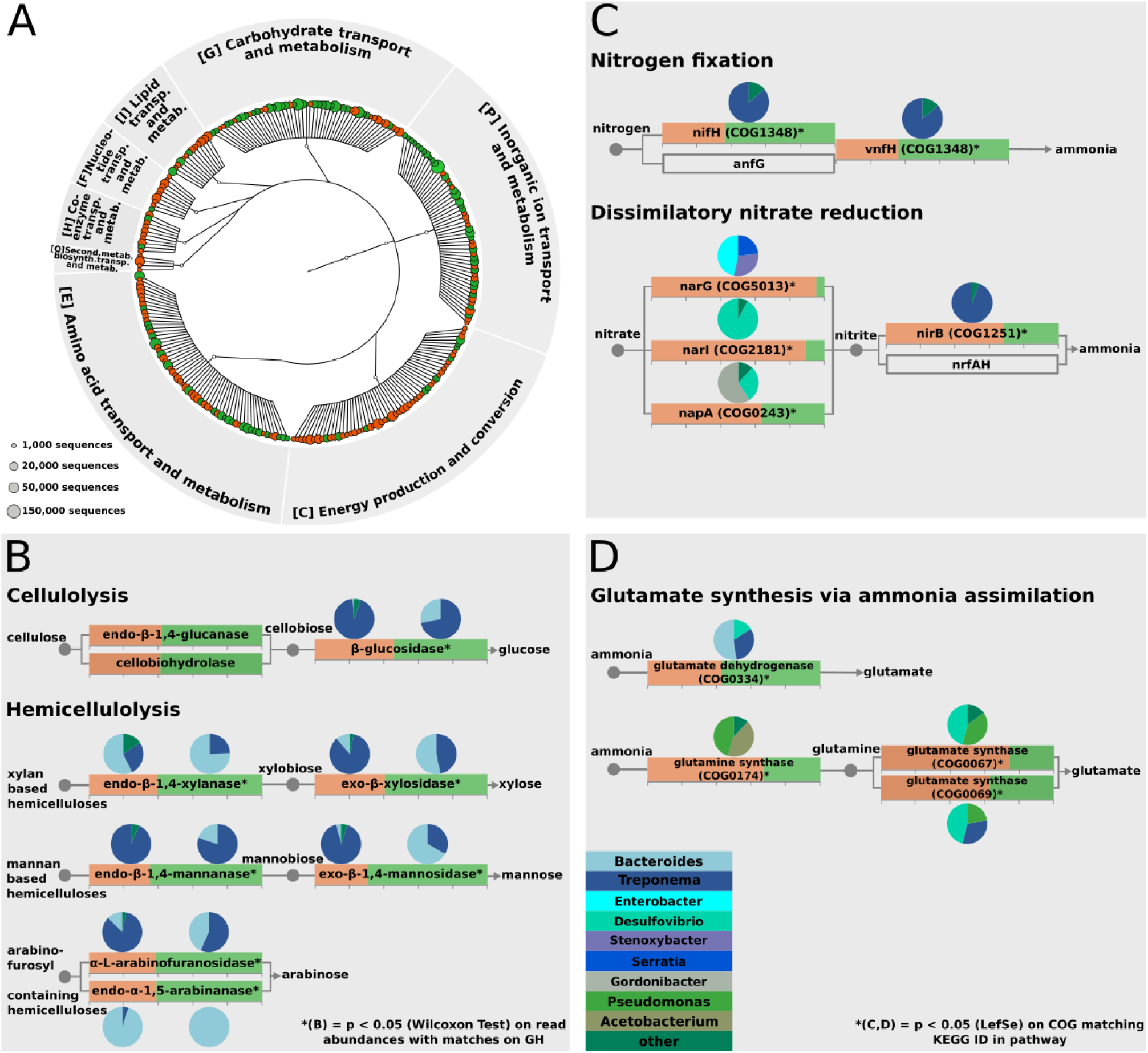
Differences in the functional content of the bacterial metagenomes of wood-dwelling and foraging species. A) Circular dendrogram/hierarchy of all COGs in the category “metabolism” over-represented in wood-dwelling species (green) or foraging species (orange). Circle size at the leafs scales with the abundance of the COG. Over-representation was detected with LefSe [57] (p < 0.05, q < 0.05, LDA > 2). A Venn diagram visualizing the total number, and the differentially abundant number of functions in each of the five pathways that constitute the category “metbolism” can be found in Supplementary Figure S7. B) Pathway analysis of cellulose and hemicellulose degradation. Colored boxes of cellulolytic or hemicellulolytic genes indicate proportion of relative abundance of sequences affiliated with wood-dwelling (green) or foraging (orange) species. C) Pathway analysis of nitrogen metabolism. Boxes for genes with functions in nitrogen metabolism indicate relative abundance in the two life types. D) Pathway analysis of glutamate synthesis. Boxes in C) and D) show relative abundance in the two life types of genes with functions in nitrogen/glutamate metabolism. Pie charts show taxonomic association of the gene. All hemicellulolytic genes were over-represented in the wood-dwelling species. Also, a nitrogenase was enriched in the wood-dwelling species, while in the foraging species, genes involved in dissimilatory nitrate reduction were over-represented.

Because wood is poor in nitrogen, termites depend on an efficient system for conserving and upgrading nitrogen [6]. In the wood-dwelling species, a potential nitrogenase ((*nif*H) COG1348)) was over-represented (Figure 4C, Table S4). Nitrogenases are key enzymes in the fixation of atmospheric nitrogen and downstream ammonia synthesis. Nitrogenase activity was mainly affiliated with members of the genus *Treponema* (Figure 4C). In contrast, in the foraging species, COGs involved in dissimilatory nitrate reduction (COG1251, COG5013, COG2181, COG0243, Figure 4C, Table S4) were over-represented. They were affiliated with a variety of different genera ranging from *Desulfovibrio* and *Gordonibacter* to *Stenoxybacter*, *Enterobacter* and *Serratia*. *Serratia* and *Enterobacter* are potential insect pathogens and contributed to the prevalence of one of the three nitrate reductases, narG (COG5013). Closer inspection of the source of these bacteria revealed that they mainly stemmed from the abnormal samples Rg2 and Rg4 that we suspected to carry a potential pathogenic infection. When we remove these samples from the analysis the increase of narG in foragers remains significant (p = 0.034).

For living on a nitrogen poor substrate, it can also be adaptive to effectively recycle nitrogen from the main waste product of the host’s amino acid metabolism, uric acid. Uric acid can be recycled through anaerobic ammonia production and downstream glutamate synthesis [6, 15, 20, 63]. In the wood-dwelling species a putative glutamate dehydrogenase (COG0334), involved in glutamate synthesis by ammonia assimilation, was over-represented. This glutamate dehydrogenase gene was mainly affiliated with members of the genera *Bacteroides*, *Treponema* and *Desulfovibrio*. In the foraging species, COGs with putative glutamine (COG0174) and glutamate synthase (COG0067, COG0069) function, were enriched (Figure 4D). These COGs were affiliated with *Desulfovibrio*, *Treponema*, *Pseudomonas* and *Acetobacterium*.

## Discussion

In this study, we assessed functional differences of termite metagenomes that underwent an evolutionary switch from wood-dwelling to foraging to identify putative contributions of the microbiome to ecological niche adaptation. To do this, we chose a set of five termite species (two foraging, three wood-dwelling species) and determined whether the functional profiles of the termite gut microbiome followed phylogeny of the host or aligned with host ecology. We hypothesized that alignment of microbiome function with termite life type is consistent with a contribution of the microbiome to termite holobiont adaptation to different ecologies. By comparing the functional content of microbiomes of different host species we focused on long-term evolutionary processes.

A potential pitfall of such an approach is that an alignment of the termite microbiome with life type-related ecology could also be caused by short-term differences between microbiomes that are merely transient. For example, microbes in the environment might differ between collection sites for the different host species. Further, ingestion of environmental microbes might lead to an association between microbiome and ecology. Similarly, differences in local food supply can lead to transient, short-term effects on the termite microbiome [55]. Consequently, such short-term differences reflect environmental differences at termite collection sites, rather than potentially adaptive, evolved differences between host-species-specific microbiomes.

For this reason, we chose to follow an approach where we control for environmental and dietary differences by acclimating all termites on the same (sterile) food source and to identical environmental conditions. We consider metagenomic patterns that persist under such highly controlled experimental conditions as robust and indicative of long-term, evolutionary acquired differences, rather than short-term imprints originating from differences in the environment or food source. It should be noted that the experimental setup poses a restriction to the number of sampled host species [33].

### Increased potential for replication in the protists of foraging termite species

In the protist metagenome of foraging species, genes involved in replication were more abundant. High replication rates are expected to be more frequently under positive selection during recolonization of the gut with protists, when the gut environment has not yet reached carrying capacity [64]. Therefore, we would like to speculate that this difference is related to the fact that *Reticulitermes* guts have to be recolonized more frequently because they molt more frequently; the intermolt periods in *Reticulitermes* are about two weeks long [49], while they average almost two months in *Cryptotermes* [48]. During molting the protists are lost and the guts have to be recolonized through proctodeal trophallaxis from nest mates [65]. However, we are aware that differences in the relative abundance of housekeeping genes like those required for replication between protist microbiomes can not be clearly disentangled from differences in average protist genome size and therefore should be interpreted with caution.

### Enrichment of genes for lignocellulose degradation in the microbiome of wood-dwelling termite species

While genes involved in replication differentiated the protist metagenomes of wood-dwelling and foraging species in our study, metabolic genes differentiated the bacterial metagenomes. Consistent with differences in their respective diets, the metagenomes of foraging and wood-dwelling species in our study differed by their potential for cellulose and hemicellulose utilization. Several GHs that have cellulolytic and hemicellulolytic function were over-represented in the metagenomes of wood-dwelling species (GH families 2, 3, 16, 43, mannosidases, xylosidases, glucanases, xylanases, Figure 4B, Table S4). A more detailed pathway analysis confirmed that hemicellulases are more abundant in the wood-dwelling species. This suggests a more pronounced role for lignocellulose degradation in the metabolism of the wood-dwelling species in our study. Accordingly, TonB dependent transporters were enriched in the microbiome of wood-dwellers. These transporters can shuttle hemicellulose and its building-blocks, in particular xylans and xylose through bacterial membranes [66, 67]. A large fraction of cellulases, hemicellulases, and putative TonB transporters were attributed to the genus *Bacteroides.* In *Bacteroides*, TonB dependent transporters are often co-localized and co-regulated with enzymes for polysaccharide degradation like hemicellulases [59, 68]. This suggests a partnership of enzymes and transporters in polysaccharide degradation. *Bacteroides* species from the human gut are also hemicellulose degraders [69], suggesting a distinctive role for the genus in hemicellulose degradation in termites as well.

The above-identified differences in functional potential between the wood-dwelling and foraging species in our study are suggestive of adaptations to utilize diets that differ in hemicellulose content. Hemicellulose content differs between wood species [70, 71]. The wood-dwelling *Cryptotermes* species in our study are mostly found in hardwood mangroves [72] where they can thrive on a bonanza food resource. The other wood-dwelling genus in our study, *Prorhinotermes,* lives in similar coastal habitats with a similar arboreal flora [73]. Hardwood is richer in hemicelluloses and the potential to use hemicelluloses is larger in the microbiome of species living on hardwood. On the other hand, *Reticulitermes* species originated in inland habitats [74], prefer soft woods like pine [75, 76] with lower hemicellulose levels, and accordingly, hemicellulolytic pathways are depleted.

### Termites with different life-types rely on different forms of nitrogen uptake and recycling

Nitrogen is scarce in a wood-based diet. As a consequence, termites need to acquire additional nitrogen from the environment. The microbiome is essential for this process. In the microbiome of wood-welling species, which feed on a uniform lignocellulose diet, a potential nitrogenase gene was enriched (nifH, COG1348). Nitrogenases are the key enzymes in the fixation of atmospheric nitrogen and downstream ammonia synthesis. This nifH was mainly affiliated with treponemes that have been shown to play an important role in nitrogen fixation before [12, 18, 19]. In contrast, the microbiome of the foraging species in our study has a higher potential to provide nitrogen to the termite holobiont by dissimilatory reduction of nitrate (Figure 4C). Nitrogen in the form of nitrate naturally occurs in soil. *R. flavipes* has been shown to acquire micro nutrients from soil [77] and to actively balance mineral uptake by food choice [78]. Therefore, it seems reasonable to assume that the microbiome of *Reticulitermes* relies on nitrogen from soil in the form of nitrate to balance the low nitrogen content of wood. The necessary nitrate reductases were found primarily in *Desulfovibrio*, *Gordonibacter* and *Stenoxybacter* that were found in association with *Reticulitermes* before and are shared between a wide range of termites [33, 79, 80].

Aside from obtaining nitrogen from the environment (atmosphere, soil), bacteria can also recycle uric acid nitrogen. All of these processes result in ammonia synthesis, the central metabolite of nitrogen metabolism. Ammonia is then further assimilated to glutamate. In the wood-dwelling species a glutamate dehydrogenase (COG0334) was over-represented. It was mainly affiliated with members of the *Bacteroides*, *Desulfovibrio* and treponemes. The foraging species seem to rely on another glutamate synthesis pathway, including glutamine (COG0174) and glutamate synthases (COG0067, COG0069). Accordingly, they were associated with a different set of bacteria including *Pseudomonas, Acetobacterium*, *Desulfovibrio,* and treponemes (Figure 4D).

### Phylogeny and ecology align with metagenome-encoded functions

Differences in the propensity for nitrogen uptake and recycling are likely to reflect differences in diet of the termite host species. Given differences in diet between the species that represent the different life types, it seems also reasonable to suggest that the changes in the repertoire of hemicellulases reflects adaptations of the microbiome to diets with different hemicellulose content. The finding that this manifested specifically in the metabolic functional repertoire, may suggest that potential selection acts in particular on metabolic functions.

Metabolic microbiome mediated adaptation to different diets can happen in two ways. First, acquisition of new microbes with adaptive functions could lead to adaptive changes of the microbiome. Second, genome evolution of microbes that are already associated with the host could lead to adaptation. Microbes that were already present before the onset of lineage specific adaptation are likely to be shared among host species. By contrast, newly acquired microbes are expected to be host lineage specific. We found that the bacterial groups that contributed most to the differentiation of metabolic functions are shared among all five host species (*Treponema*, *Bacteroides*, *Desulfovibrio*, *Dysgomonas, Gordonibacter, Pseudomonas*, Table S4, Figure S3). This supports that genome evolution of microbes that were already associated with the host contributed to potential adaptation in our model system.

## Conclusion

We applied metagenomic sequencing of gut microbiomes from a controlled experimental setup to assess a putative contribution of the microbiome to host ecological adaptation that accompanies the evolutionary switch from wood dwelling to foraging life types. We found that the overall pattern of microbiome variation reflected a phylogenetic signal. Interestingly, however, specific functions of the microbiome aligned with the underlying host ecology. The specific ecology related differences in microbiome function led us to hypothesize that the microbiome contributed to dietary adaptations, namely different hemicellulose and nitrogen contents. This hypothesis can now be tested, assessing host fitness under different dietary conditions. Such experiments will be crucial to disentangle adaptive functional changes from selectively neutral functional turnover or side effects of other adaptations.

## Experimental Procedures

### Termite samples

All termites were collected from typical natural habitats (see [33]). They were kept under constant conditions (27°C, 70% humidity) on autoclaved *Pinus radiata* wood from the same source for at least six weeks prior to the experiment. The feeding of *Pinus* represents a natural or near natural treatment; *Pinus* is a natural food source of *P. simplex* and *Reticulitermes. Cryptotermes* growth and behavior on *Pinus* recapitulates that on natural substrate [72]. The time of the acclimation period was chosen to lie well beyond the gut passage time of 24 h in lower termites [81, 82] and following Huang et al. [83], who showed that six weeks are sufficient for the microbiota to adjust to a new diet. That way, all excretable material like remaining food, transient microbes taken up from the environment that have no mechanisms to persist in the gut, and microbial DNA taken up before the experiment was made sure to be excreted. The samples were identical to those analysed in our previous study, [33] where detailed information about animal collection, keeping, and cytochrome oxidase II based species identification and a phylogeny can be found.

### DNA extraction and Shotgun sequencing

DNA was extracted from a pool of three worker guts per colony using bead beating, chloroform extraction and isopropanol precipitation (see Supplementary material and methods file S7). Each of the 29 colony samples went through independent metagenomic shotgun library preparation and sequencing on an Illumina HiSeq platform (150 bp paired end reads).

### Analysis

We employed a double filtering strategy to remove host DNA from our analysis. First, sequences were removed that mapped to available host genomes from *C. secundus* [84] and transcriptomes from *P. simplex* [85] and *R. flavipes*, provided by the 1KITE consortium (www.1kite.org, BioSample SAMN04005235) using BBMap [86] (for detailed workflow and more detailed information about used genomes and transcriptomes see Supplementary Figure S2 and file S8). Of note, the sequences were not assembled, but individual reads were directly annotated. In a second step we used taxonomic and functional annotations with Megan6 [87] to retrieve only sequences that could be unambiguously assigned to either bacteria or protists. In order to compare the bacterial and protist data sets of all samples, they were rarefied to the number of sequences in the sample with lowest coverage, resulting in 1,386,882 and 2,781 sequences per sample, respectively. Sample Cs4 was excluded from the analysis for insufficient sequence coverage (974,176 sequences), so was Cs5 from the protist data. Sample Ps5 did not pass the analysis pipeline and was also excluded.

Functional annotation with the eggNOG database resulted in the highest number of annotated sequences (21,215,480 annotated sequences in total) and was chosen for further functional analysis. Bray-Curtis distances of functional abundances were clustered with the pvClust package in R [88]. Multivariate modeling was performed via RDA (Redundancy Analysis) and AICs as well as values for the proportion of variance explained were derived with the model selection tool ordistep and ordiR2step, as implemented in the R vegan package [89]. Models were compared to the null-model via ANOVA. To identify over-represented functions associated with the two termite life types, a Linear Discriminant Analysis (LDR) was performed using LEfSe [57] and visualized using graphlan [90]. Pathway analysis of CAZy GHs was performed by blasting bacterial reads of all samples against the full CAZy protein database, using Diamond [91]. GH abundance was estimated by counting reads with matches on proteins with cellulolytic and hemicellulolytic functions [92]. Pathway analysis of the nitrogen metabolism was performed by searching COG IDs corresponding to the KEGG IDs among the over-represented COGs from the LEfSe analysis. A detailed workflow for full reproducibility can be found in Supplementary Figure S2 and file S10 and S11.

## Supporting information

supplementary figures and analysis workflow

## Acknowledgments

We thank Jan Sobotnik for kindly providing *P. simplex* colonies, as well as Charles Darwin University (Australia), especially S. Garnett and the Horticulture and Aquaculture team for providing logistic support to collect *C. secundus*. The Parks and Wildlife Comission, Northern Territory, Department of the Environment, Water, Heritage and the Arts gave permission to collect (Permit number 59044) and export (Permit PWS2016-AU-001559) the termites. This work was funded by DFG grants STA1154/2-1 and KO1895/16-1. We thank Craig Michell and the KAUST Bioscience Core Lab for sequence library generation and sequencing. This study was supported by King Abdullah University of Science and Technology (KAUST) and the High Performance and Cloud Computing Group at the Zentrum fuer Datenverarbeitung of the University of Tuebingen, the state of Baden-Wuerttemberg through bwHPC and the German Research Foundation (DFG) through grant no INST 37/935-1 FUGG. We thank Karen Meusemann and the 1KITE consortium, in particular the 1KITE Blattodea group, Alexander Donath, Lars Podsiadlowski, Bernhard Misof, Xin Zhou for granting access of the transcriptome data of *R. santonsensis* syn. *R. flavipes*.

## Declarations

### Ethics approval and consent to participate

Not applicable.

### Availability of data and material

The raw data has been uploaded to the ncbi short-read archive (BioProject ID PRJNA509211, Accession: SAMN10573992 – SAMN10574019). Supporting information and analysis workflows are included in the supplementary files in this article.

### Author contributions

FS, JK, LW designed the experiment. JK, FD provided study organisms. LW performed the experiments, CV generated the sequence data, LW and FS analyzed the data. LW, FS, JK, CV, and FD wrote the manuscript. All authors read and approved the final manuscript.

### Consent for publication

Not applicable.

