## supplementary figures and analysis workflow for "Ecological specificity of the metagenome in a set of lower termite species supports contribution of the microbiome to adaptation of the host"

### Supplementary tables, figures and information

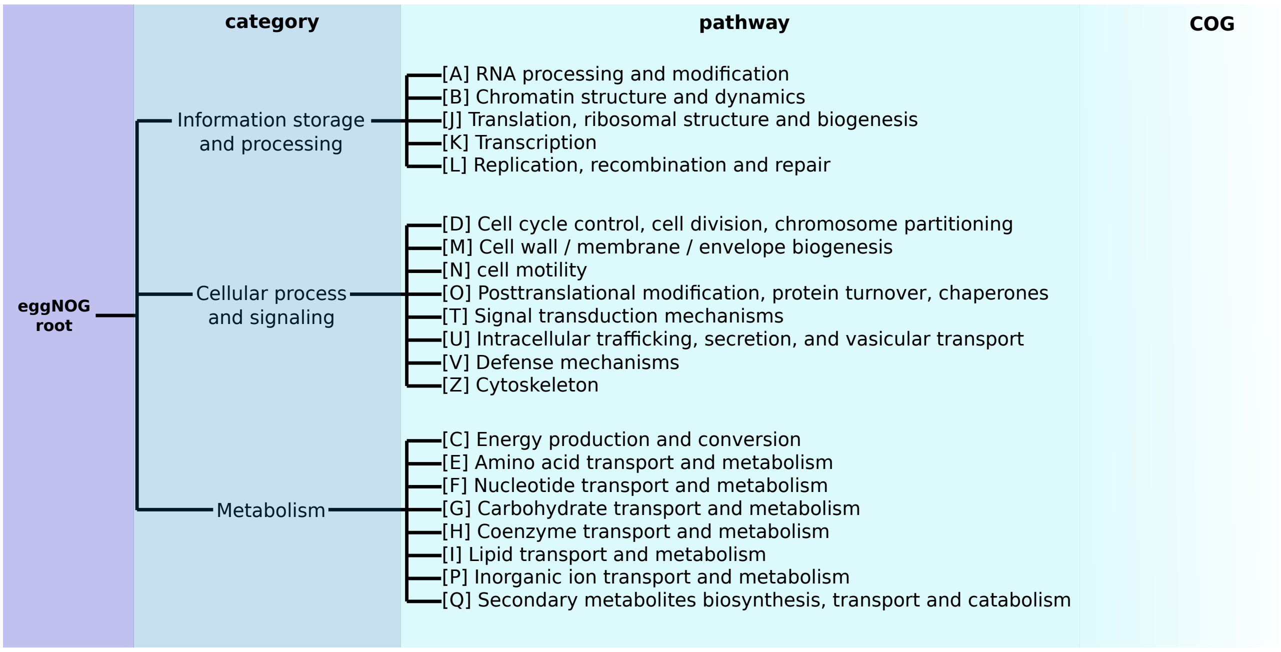


**Figure S1: eggNOG annotation structure.** The eggNOG annotation [1] is divided into three categories and 21 pathways into which the COGs are grouped.

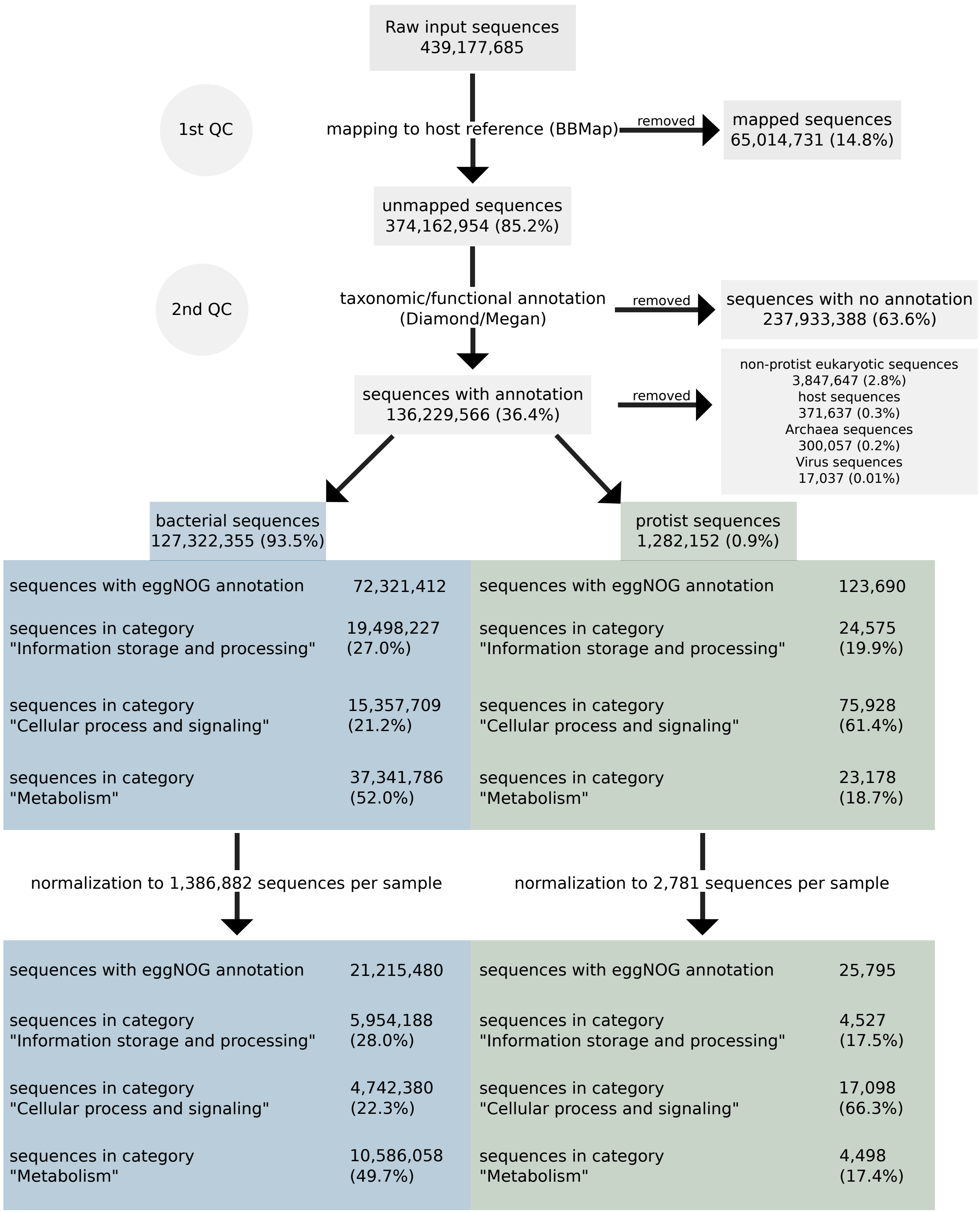


**Figure S2: Analysis workflow.** Of note, sequences were not assembled, since resulting contigs were not significantly longer than individual reads, likely due to the complexity of the microbiomes (data not shown). As a first filtering step, raw sequences were mapped against a host reference, using BBMap (version 37.02, [2]), with standard settings. *C.*

*secundus* and *C. domesticus* samples were mapped against the *C. secundus* [3] genome. *P. simplex* samples were mapped against the *P. simplex* transcriptome [4], *R. flavipes* and *R. grassei* samples against the *R. santonensis* transcriptome (provided by the 1KITE consortium (www.1kite.org, BioSample SAMN04005235 (for details on sequencing and removal of non-termite contaminants and cross-contaminants from the 1kite data, see [5]). Only sequences that did not match the host reference ("unmapped sequences") were used for further analysis. Taxonomic and functional annotation of unmapped sequences was performed using Diamond (version 0.8.37.99, [6]) and Megan (MEGAN Community Edition (version 6.7.18, [7]). As a second filtering step, only sequences with taxonomic classification "bacteria" or "parabasalia" were exported into two separate files for each sample. Therefore, contaminants such as left over host sequences, archaea, or virus sequences were removed from the datasets. In order to compare the bacterial and protist metagenomes of all samples, they were normalized to 1,386,882 and 2,781 sequences per sample, respectively. See also Supplementary file S10 below for detailed analysis steps.

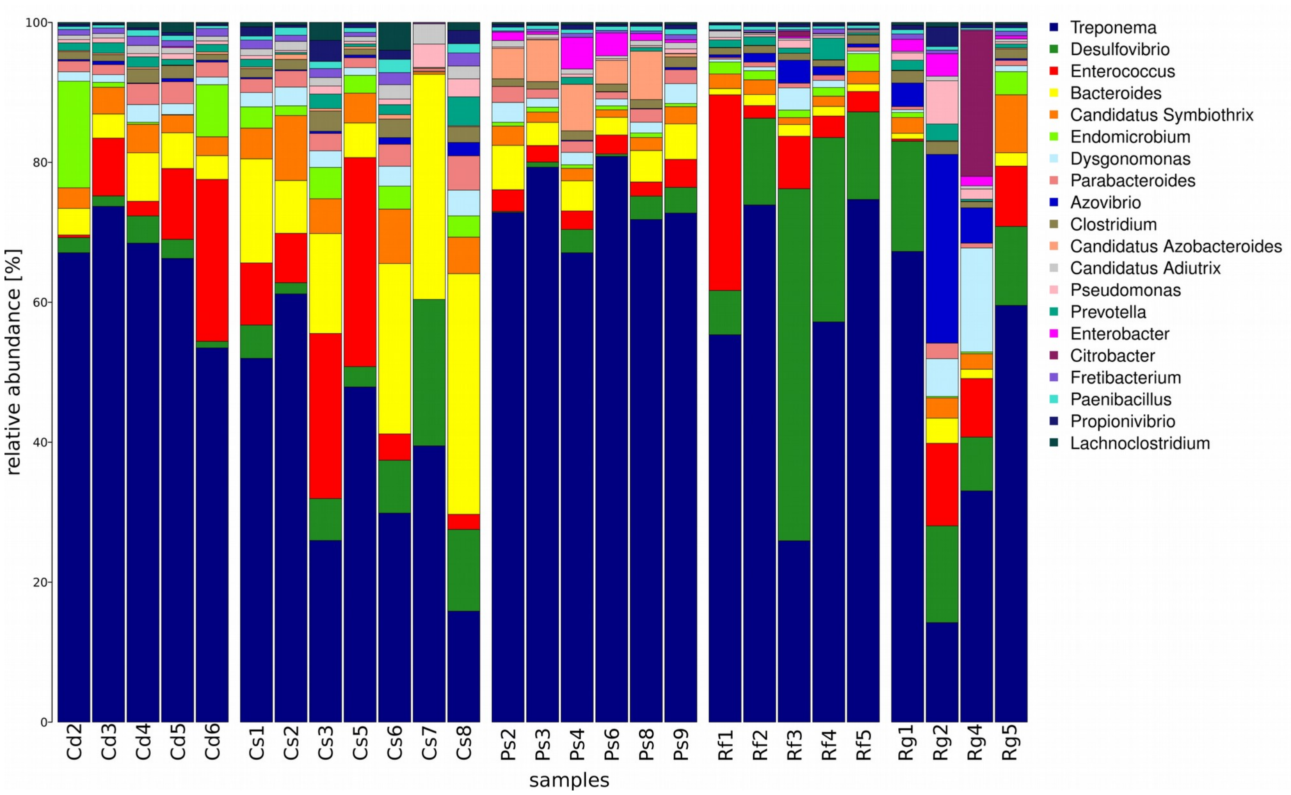


**Figure S3: Frequency of bacterial genera.** Cd = colony replicates of *C. domesticus*; Cs = colony replicates of *C. secundus*; Ps = colony replicates of *P. simplex*; Rf = colony replicates of *R. flavipes*; Rg and  = colony replicates of *R. grassei*. Shown are only the 20 bacterial genera with the highest sequence coverage. *Treponema* was the most abundant genus (14.2% (Rg2) – 80.9% (Ps6)), followed by *Desulfovibrio* (0.1% (Ps2) - 50.3% (Rf3)), *Enterococcus* (0% (Cs7) – 29.9% (Cs%)) and *Bacteroides* (0.8% (Rf1) – 34.4% (Cs8)). Like already observed in a previous study with the same samples [8], Rg2 and Rg4 showed an unusual community structure.

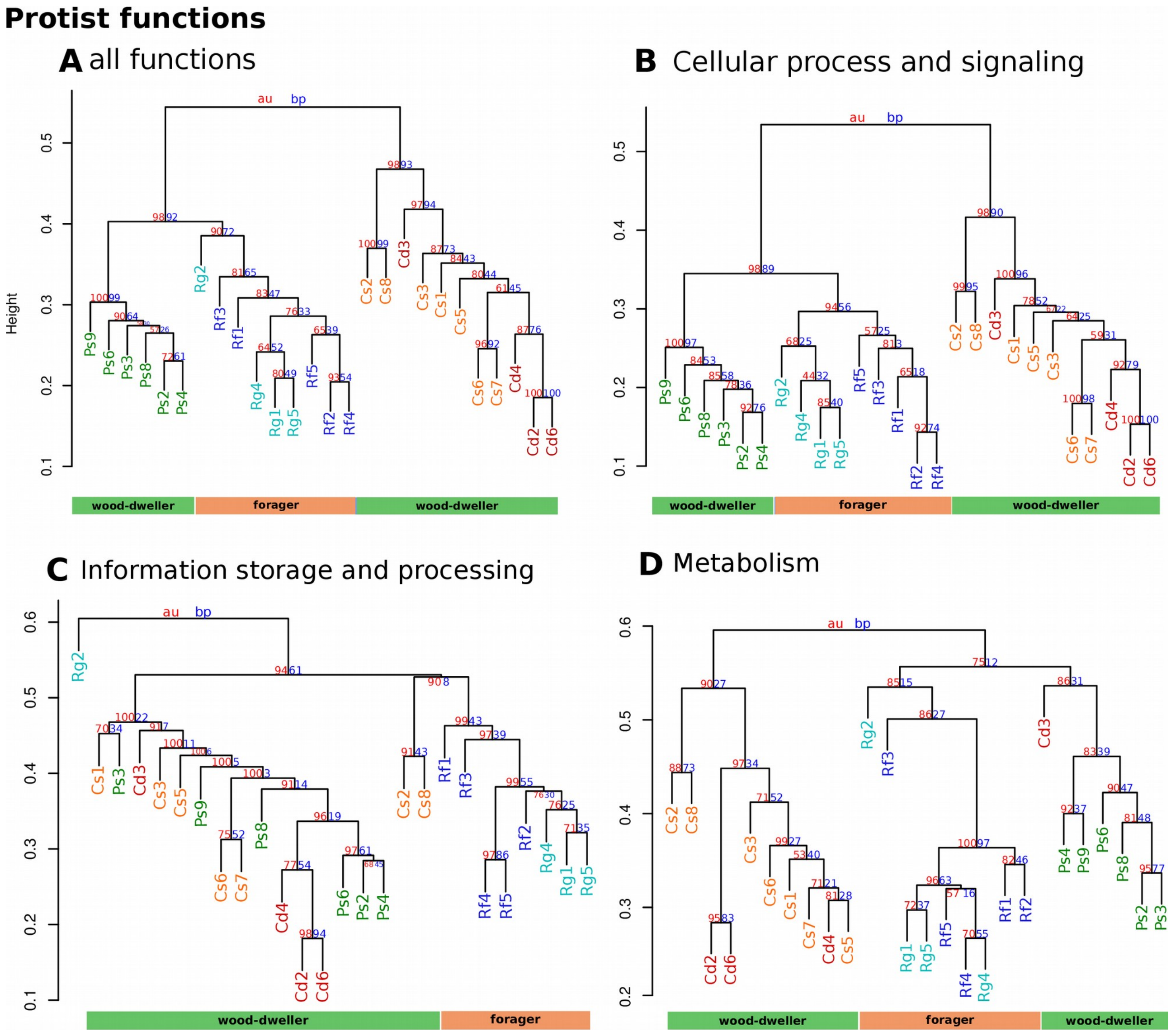


**Figure S4: Cluster dendrograms of the functional profiles of the protist community.** Community distances are based on Bray-Curtis Dissimilarities. A) all functions (25,795 sequences). B) category “cellular process and signaling” (17,098 sequences). C) category “information storage and processing” (4,527 sequences). D) category “metabolism” (4,498 sequences). Cd (red) *C. domesticus* colonies; Cs (orange) *C. secundus* colonies; Ps (green) *P. simplex* colonies; Rf (blue) *R. flavipes* colonies; Rg (lightblue) *R. grassei* colonies. All functional profiles show a strong phylogenetic imprint. However, functions involved in “information storage and processing” were more similar between hosts of the same ecological life type.

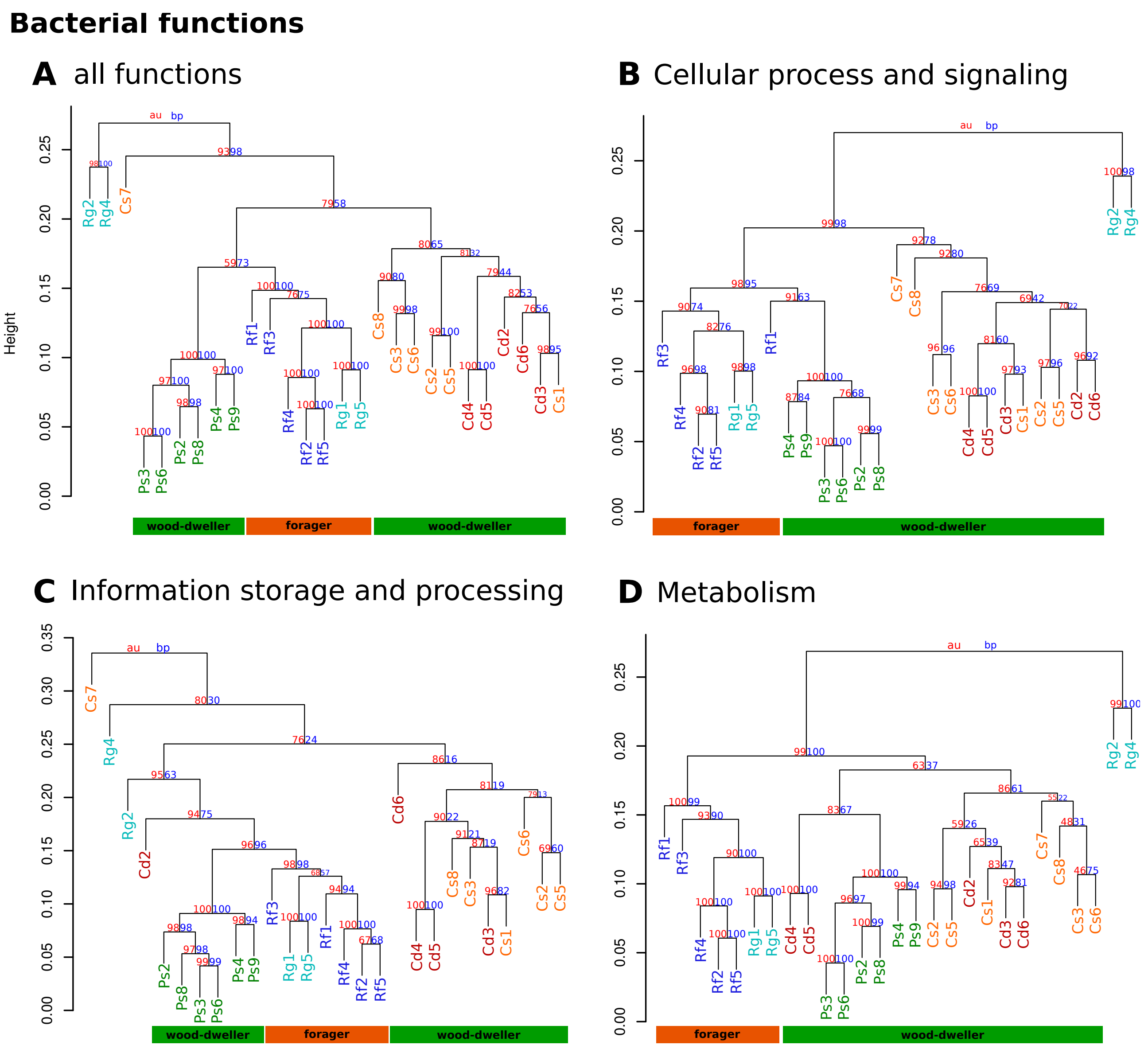


**Figure S5: Cluster dendrograms of the functional profiles of the bacterial community.** Community distances are based on Bray-Curtis Dissimilarities. A) all functions (21,215,480 sequences). B) category “cellular process and signaling” (4,742,380 sequences). C) category “information storage and processing” (5,954,188 sequences). D) category “metabolism” (10,586,058 sequences). Cd (red) *C. domesticus* colonies; Cs (orange) *C. secundus* colonies; Ps (green) *P. simplex* colonies; Rf (blue) *R. flavipes* colonies; Rg (lightblue) *R. grassei* colonies. Similar to the protist set, bacterial functional metagenome showed a strong phylogenetic imprint. However, the metabolic metagenomes clustered according to host ecology (life type).

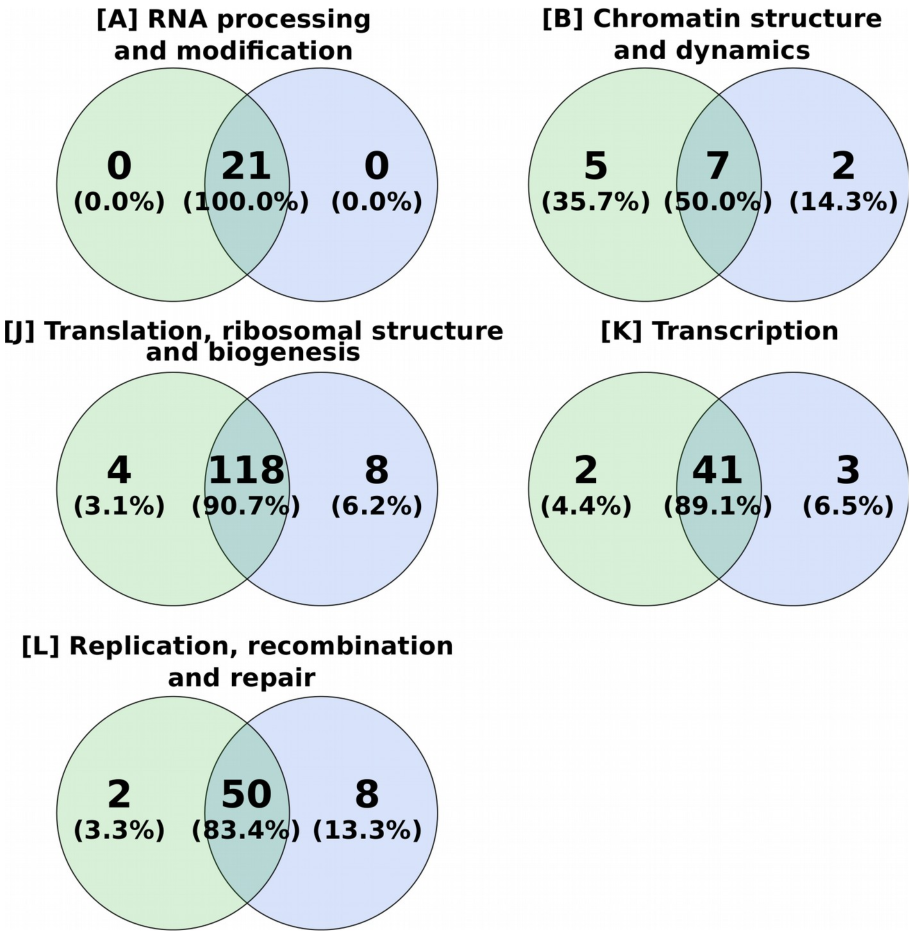


**Figure S6: Venn diagram of COGs in the pathways of the eggNOG category "information storage and processing" in the protist metagenoms.** Shown are numbers and percentages of  over-represented and shared COGs found in the protist metagenomes of wood-dwelling (green) and foraging (orange) termite species.

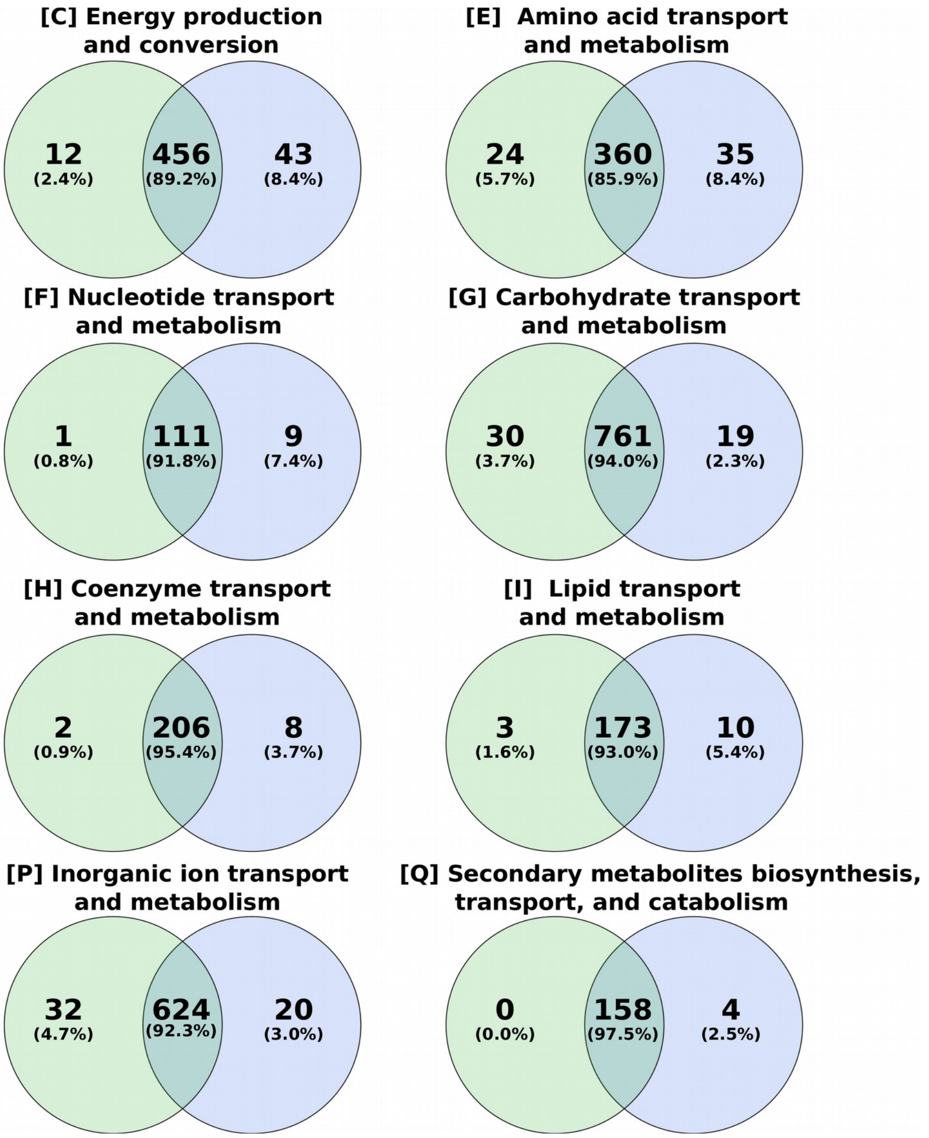


**Figure S7: Venn diagram of COGs in the pathways of the eggNOG category "metabolism" in the bacterial metagenoms.** Shown are numbers and percentages of over-represented and shared COGs found in the bacterial metagenomes of wood-dwelling (green) and foraging (orange) termite species.

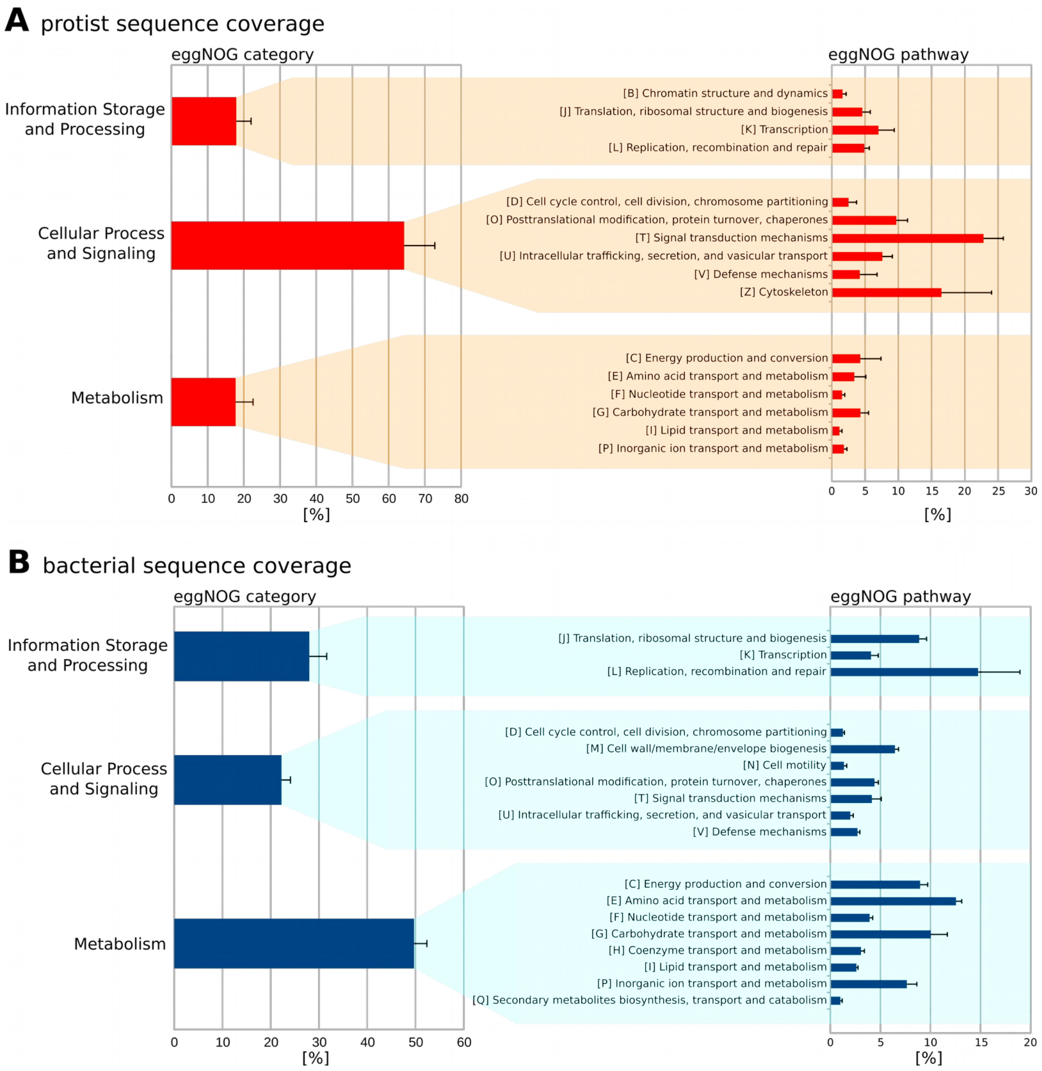


**Figure S9: Metagenome sequence coverage of functional categories.** A) protist sequences. B) bacterial sequences. Pathways with less than 1% coverage were removed for clarity. Background highlighting represents grouping of pathways into categories according to the eggNOG hierarchy (Figure S1). More than half (64.3%) of the protist metagenome were assigned to “cellular process and signaling”. In the bacterial metagenome sequences with annotations related to metabolic functions comprised the major group (49.7%). Error bars represent standard deviation across all samples.

<u>Functional potential of protist and bacterial metagenomes</u>

The largest group of functions encoded by the protist metagenome were related to signaling (Figure S9A). In particular, ABC-transporters were abundant. ABC-transporters can transport a large variety of substrates in and out of the cell [9]. In parasitic protists, ABC transporters are essential for drug resistance and nutrient salvation [10]. They also play an important role in several insect-microbe-symbioses, in particular, when metabolic pathways are split between partners and intermediate metabolites have to be shuttled between organisms [11]. Therefore, we speculate here that shuttling of molecules is important to protists in the termite gut because they occupy a mediator position between their intracellular endosymbiotic bacteria and the termite host. Metabolites and other molecules cannot be directly exchanged between termites and intracellular bacteria, instead transport through the protist is required. This in return requires transmembrane transporters and might explain their abundance.

In contrast to the prevalence of transporters in protists, the largest fraction of bacterial sequences had metabolic functions (Figure S9B). The most common metabolic category was 'amino acid metabolism' including nitrogen metabolism. The microbiome plays a central role in nitrogen uptake and recycling because the host's primary food source, wood, is poor in nitrogen [12–14]. The second most common pathway was carbohydrate metabolism. In this pathway, we identified a total of 99 glycoside hydrolases from 31 different families (Table S5). These are the primary enzymes responsible for lignocellulose degradation. Thus, our results support a growing body of evidence that bacteria play a fundamental role in lignocellulose degradation that was formerly mainly attributed to protists in lower termites [15–18].

**Table S1: Sample Overview.** Number of sequences before and after quality filtering, as well as taxonomic and functional annotation statistics.

**Table S2: Transposase abundance in the bacterial dataset in the pathway “replication, recombination and repair”.** COGs annotated as transposase are marked in color.

**Table S3: Complete list of results of Linear Discriminant Analysis (LDA) of protist dataset.** COGs were grouped into categories and color coded based on their function.

**Table S4: Complete list of results of Linear Discriminant Analysis (LDA) of bacterial dataset.** COGs were grouped into categories and color coded based on their function.

**Table S5: Complete list of bacterial COGs found within the pathway 'carbohydrate transport and metabolism'**. COGs identified as glycoside hydrolase (GH) marked in yellow.

**Supplement S10. Metagenomic Shotgun Data Analysis Workflow**

All computing steps in this script were performed on a High Performance Computing Cluster (bwForCluster BinAC, Eberhard Karls University of Tuebingen, High Performance and Cloud Computing Group at the Zentrum fuer Datenverarbeitung of the University of Tuebingen, the state of Baden-Wuerttemberg through bwHPC and the German Research Foundation (DFG) through grant no INST 37/935-1 FUGG).

1.) Please note that the sequences were not assembled (see Material and Methods main text), but directly annotated using the following workflow.

1.) Mapping raw input sequences against reference genomes with BBMap (version 37.02, [2]).

Sequences of *Cryptotermes secundus* and *Cryptotermes domesticus* were mapped against the *C. secundus* genome [3], sequences of *Prorhinotermes simplex* against the *P.*

*simplex* transcriptome ([4], NCBI Bioproject ID: 219597, Assembly version GASE02000000) and sequences of *Reticulitermes flavipes* and *Reticulitermes grassei* against the *R. santonensis* (syn. *R. flavipes* [19]) transcriptome provided by the 1KITE consortium (www.1kite.org, BioSample SAMN04005235 (for details on sequencing and removal of non-termite contaminants and cross-contaminants from the 1kite data, see [5]).

```
>module load devel/java_jdk/1.8.0u112 #load required Java
environment
>cd path/to/directory #change to working directory
>/path/to/software/bbmap.sh
ref=reference_genome_or_transcriptome.fa
in1=Sample1_Read1.fastq.gz in2=Sample1_Read2.fastq.gz
outm1=Sample1_Read1_mapped.fastq.gz
outm2=Sample1_Read2_mapped.fastq.gz
outu1=Sample_Read1_unmapped.fastq.gz
outu2=Sample1_Read2_unmapped.fastq.gz #run BBMap with standard
settings, do for all samples
```

This will produce four output files: Sample_Read1_mapped.fastq.gz, Sample_Read1_unmapped.fastq.gz,Sample_Read2_mapped.fastq.gz, Sample_Read2_unmapped.fastq.gz. Only “unmapped” sequences, that did not match the host reference, were used for further analysis.

2.) SYNTAX for Diamond and Megan6

This script is following the Syntax tutorial as provided by the BinAC HPC. For Diamond (version 0.8.37.99, [6]), the non-redundant NCBI database was used as reference, for functional classification with Megan6 (MEGAN Community Edition (version 6.7.18, [7]), the mapping files prot_acc2tax-May2017.abin.gz, acc2eggnog-Oct2016X.abin.gz, acc2interpro-Nov2016XX.abin and acc2seed-May2015XX.abin (download details below) were used.

```
>wget ftp://ftp.ncbi.nih.gov/blast/db/FASTA/nr.gz #download NCBI
nr.gz database
> diamond makedb --in nr.gz --db nr #build Diamond index for the
nr database, will generate a new file called nr.dmnd
```

The mapping files for Megan6 were downloaded from the Megan6 download page (http://ab.inf.uni-tuebingen.de/data/software/megan6/download/welcome.html)

```
>mkdir path/to/wanted/directory/00fastq #put unmapped BBMap output
files in here
>mkdir path/to/wanted/directory/10daa #will contain daa files
generated by Diamond
>mkdir path/to/wanted/directory/20rma #will contain rma files
generated by Megan6
```

For each file fastq.gz in the 00fastq directory, run Diamond as follows:

```
>module load bio/diamond/0.8.37 #load diamond environment
>module load bio/megan/6.7.18 #load Megan6 environment
>cd path/to/working/directory #change to working directory
>diamond blastx --query 00fastq/Sample1_Read1_unmapped.fastq.gz --
db nr.dmnd --daa 10daa/Sample1_Read1_unmapped.daa #run Diamond,
will generate a daa Diamond output file in the /10daa directory
```

For each daa file in your 10daa directory, run Megan's daa2rma as follows:

```
>module load devel/java_jdk/1.8.0u112 #load required Java
environment
>module load bio/diamond/0.8.37 #load Diamond environment
>module load bio/megan/6.7.18 #load Megan6 environment
>cd path/to/working/directory # change to working directory
>daa2rma -p -ps 1 -i 10daa/Sample1_Read1_unmapped.daa
10daa/Sample1_Read2_unmapped.daa -o 20rma/Sample1_p_unmapped.rma -
a2t prot_acc2tax-May2017.abin -a2eggnog acc2eggnog-Oct2016X.abin -
a2seed acc2seed-May2015XX.abin -a2interpro2go acc2interpro-
Nov2016XX.abin -fun EGGNOG SEED INTERPRO2GO #the -p function is
used for a paired read input and will generate a merged rma6
output file in the /20rma directory. This file can be used as
input for the Megan6 graphical user interface.
```

Further analysis using the Megan6 graphical user interface was performed locally. Each rma6 file of each sample was imported in Megan6 individually. Then bacterial and protist reads were extracted into separate files using the "Extract reads.." function in Megan6 (see also Material and Method section). To compare the bacterial and protist datasets of all samples, they were imported into Megan6 using the "Compare..." function. During this step, the bacterial and protist datasets were normalized to 1,386,882 and 2,781 reads respectively.

3.) PvClust Cluster Dendrograms

Cluster dendrograms were generated in Rstudio (version 3.3.1, [20]) using the PvClust package and an external script (https://github.com/hallamlab/mp_tutorial/blob/master/taxonomic_analysis/code/mp_tutorial_taxonomic_analysis.R) to add the Bray-Curtis dissimilarity index. Functional abundance tables were extracted from Megan6 using the “Export...” function.

In Rstudio, execute:

```
>library(pvclust)
>source('path/to/working/directory/pvclust_bcdist.R')   #source  to
downloaded R script
>setwd("/path/to/working/directory") #set working directory
>read.csv("FunctionalAbundance.csv",header=TRUE,sep=",",row.names
= 1)→dat #read in input
>clust_dat     <-     pvclust(dat,     method.hclust="average",
method.dist="bray-curtis", n=1000) #use pvclust
>plot(clust_dat,cex=0.8) #plot
>pdf(“clust_dat.pdf") #export as pdf
>plot(clust_dat,cex=0.8)
>dev.off()
```

4.) RDA and model selection

Functional abundance tables were exported from Megan6 using “Export...” > “Text (CSV) format...” > “eggnogPath_to_count”. Bacterial functional abundances were rarefied to at least 1,000 sequences per COG, protist functional abundances to at least 10 sequences per COG. Functional abundances were subsequently transformed using Hellinger transformation. RDA and ordistep/ordiR2step for model selection were performed with the vegan package in R [21] and compared with ANOVA using the following commands:

```
#set Nullmodel
>rda0 <- rda(functional_abundance_table ~ 1, data = metadata)
#set model with 1 explanatory variable
>rda2  <-  rda(functional_abundance_table  ~  host_family,data  =
metadata)
>rda3  <-  rda(functional_abundance_table  ~  host_lifetype  ,data  =
metadata)
#set model with both explanatory variables for modelselection
>rda1   <-   rda(functional_abundance_table   ~   host_family   +
host_lifetype ,data = metadata)
#model selection via ordistep
>model_selection <- ordistep(rda0, scope = formula(rda1))
#model selection via ordiR2step
```

```
model_selection_inf <- ordiR2step(rda0_inf, scope = formula(rda1_inf), trace = TRUE, permutations = how(nperm = 499), Pin = 0.05, R2scope = FALSE)
#comparison against Nullmodel
>anova.cca(rda0,rda2)
>anova.cca(rda0,rda3)
```

Removing outlier samples from the datasets (outlier samples were Rg2, Cs2 and Cs8 in the category “information storage and processing” of the protist functional set and Rg2, Rg4 in the category “metabolism” in the bacterial set, and Rg2, Rg4 and Cs7 in all bacterial functions) did not change significant results.

5.) LEfSe

Functional abundance tables were exported from Megan6 using the “Export...” > “Text (CSV) format...” > “eggnogPath_to_count”. To use this file as input for LEfse [22], a row with the “class” “wooddweller” or “forager” was added above the sample names.

```
>python format_input.py FunctionalAbundance.csv FunctionalAbundance.lefse.in -c 1 -s -1 -u 2 -o 1000000 #will create the .in file
>python run_lefse.py FunctionalAbundance.lefse.in FunctionalAbundance.lefse.res #will create the .res file
```

6.) Circular dendrogram of LEfse results using GraPhlAn (Asnicar *et al*., 2015)

Only significant overrespresented fuctions (LDA > 2.0, q-value < 0.05) were used as input.

```
>python export2graphlan.py -i FunctionalAbundance.csv -o FunctionalAbundance.lefse.res -t tree.txt -a annot-txt --title "Functional Abundance" --external_annotations 4 --fname_row 0 --skip_rows 1
>python graphlan_annotate.py --annot test_annot.txt test_tree.txt test_outtree.txt
>python graphlan.py --dpi 150 test_outtree.txt test_outimg.png --external_legends
```

7.) CAZy pathway analysis.

Full CAZy reference database was downloaded from http://csbl.bmb.uga.edu/dbCAN/index.php (newest version CAZyDB.07202017.fa). Bacterial reads were blasted against the reference using Diamond:

```
>diamond makedb --in CAZyDB.07202017.fa -d cazy_full
>diamond blastx -d cazy_full -q bacteria_sampleXY.fasta -o matches.m8 -k 1 -e 0.00001 --max-hsps 1
```

Read matches for GHs of interest* were counted by piping the wc -l function after a grep search command.

*Read matches for GHs of interest: cellulose:
endo-ß-1,4-glucanase (cellulase): GH5,8
cellobiohydrolase: GH6,7,9,48,74
ß-glucosidase: GH1,3,4,5,9,13,17,30,31,63,65,97,116,122,133
hemicellulose (see [23]):
xylan:
endo-ß-1,4-xylanase: GH5,8,10,11,43
exo-ß-xylosidase: GH3,39,43,52,54
mannan:
endo-ß-1,4-mannanase: GH5,26
exo-ß-1,4-mannosidase: GH1,2,5
arabinofuranosyl containing hemicellulose:
alpha-L-arabinofuranosidase: GH3,43,51,54,62
endo-alpha-1,5-arabinanase: GH43

**S11. Supplementary Material and Methods**

DNA extraction

Entire guts of three workers per colony were extracted and immediately transferred to 100 µl of CTAB solution (0.75 M NaCl, 0.05 M Tris, 0.01 M EDTA, 2% CTAB). Guts were homogenized (zirconia and glass beats, 3 min at 25*1/s), using a Tissue Lyser II (Qiagen). Additional 400 µl of CTAB solution were added and samples were incubated for 1h at 65°C and 800 rpm on a Thermomixer Comfort (Eppendorf). After addition of 2 µl Proteinase K (Thermoscientific, concentration: ~20 mg/mL) samples were incubated for 2h at 55°C and 800 rpm. The proteinase K reaction was terminated by heating to 98°C for 15 min. DNA was extracted with 500 µl of chloroform:isoamylic alcohol (24:1) followed by centrifugation for 15 min at 10000 rpm. DNA, was precipitated with 325 µl of ice cold isopropanol and overnight incubation at -20°C. Samples were washed with 300 µl of 100% ethanol and twice with 300 µl 70% ethanol, centrifuging for 15 min at 4°C at 14.000 rpm between each washing step. The DNA pellet was air dried and resuspended in 50µl of water.
